# Transcriptional profiling of *Impatiens walleriana* genes through different stages of downy mildew infection reveals novel genes involved in disease susceptibility

**DOI:** 10.1101/622480

**Authors:** Stephanie Suarez, Zunaira Afzal Naveed, Gul Shad Ali

**Affiliations:** Tropical Research and Education Center, University of Florida, Homestead, Florida 33031, USA; Department of Plant Pathology, University of Florida, Gainesville, Florida 32611, USA; Mid-Florida Research and Education Center, Department of Plant Pathology, University of Florida, Apopka, Florida 32703, USA

## Abstract

Impatiens downy mildew is a highly destructive disease of *Impatiens walleriana*, and economically important bedding ornamental crop. This disease is caused by a recently emerged pathogen *Plasmopara obducens*. Since both the host and pathogen are relatively less studied, there are only a few genomic resources available for both *I. walleriana* and *P. obducens*. In this study, we have analyzed transcriptional changes in *I. walleriana* in response to *P. obducens* infection during different stages of disease development. Our main goal was to identify candidate genes that may be involved in *I. walleriana* susceptibility to *P. obducens*. Since the genome of *I. walleriana* is not available publicly, we constructed and optimized a *de novo* transcriptome assembly consisting of 73,022 transcripts. Differential expression analysis based on this optimized *de novo* transcriptome assembly revealed 3,000 to 4,500 differentially expressed transcripts (DETs) at 0 hr, 12 hr, 48 hr, 120 hr, and 240 hr time points after infection. Functional annotation of these DETs revealed that numerous plant stress responsive genes are activated and deactivated throughout the infection cycle. Genes in the calcium signaling pathways, receptor-like kinases (RLKs) including 10 disease resistance associated RLK transcripts, powdery mildew resistance genes (*MLO*), and many other plant stress related genes were predominantly differentially expressed in *I. walleriana* in response to *P. obducens*. Analyses reported here provides molecular insights into the disease susceptibility mechanism of the Impatiens downy mildew, and lays out a strong foundation for future studies aimed at improving downy mildew resistance in *I. walleriana*.

## Introduction

Downy mildews are notoriously destructive diseases causing devastating losses on both agronomic and ornamental crops worldwide. With reports on wild impatiens dating back to 1877 (Farr et al. 2016), the first report of impatiens downy mildew (IDM) on cultivated *I. walleriana* in the United States did not occur until 2004 (Lane et al. 2005). The fatal consequences of this disease on impatiens production have gained worldwide news coverage, as impatiens was previously the top annual bedding plant in the floriculture industry. Specifically, production in Florida averaged $20 million flats annually before IDM was first reported in the state. Since its emergence, gardening impatiens has been replaced by landscape alternatives such as begonias and the IDM-resistant New Guinea impatiens (*I. hawkeri*).

Because this pathogen is a newly emerging one, there is little known about *P. obducens*. There are few genomic resources available in regard to both I. walleriana and *P. Obducens*. Recently, the first de novo assembly of *P. obducens* resulted in a 202-Mb genome (Salgado-Salazar et al. 2015). Polymorphic SSR marker candidates were discovered and will hopefully serve as an exciting new source as the ongoing molecular studies of this pathogen continues. While there is much to learn about the genetics of *P. obducens*, recent research available regarding *I. walleriana* was published identifying disease resistance genes as well as SSRs and SNPs sites which may aid in breeding impatiens for resistance to IDM (Bhattarai et al. 2018). There is strong consumer demand for IDM-resistant *I. walleriana* cultivars. In addition to studying the host and its pathogen separately, understanding the plant defenses in the host-pathogen interaction during infection is critical in order to introduce disease resistance in susceptible host cultivars.

Pattern recognition receptors (PRRs) are the first layer of plant defense in the innate immune system of plants. Localized to host cell membranes, they recognize pathogen structures or molecules known as pathogen-associated molecular pattern (PAMP) and in turn, activate PAMP-Triggered Immunity (PTI). PTI is a non-specific defense mechanism allowing for recognition of both non-pathogenic and pathogen organisms, and thus combatting pathogen invasion and further colonization. A second layer of immunity, effector-triggered immunity (ETI) may be activated if a pathogen survives PTI. ETI depends on pathogen effectors, secreted in host cells by the pathogen, resulting in programmed cell death at the site of infection, or a hypersensitive response (HR) in an incompatible reaction.

HR is prompted by host resistance (R) genes, which are divided into five classes: (i) the CNL class comprising of R genes encoding a protein with at least one N-terminal coiled-coil, a nucleotide binding site and a leucine-rich repeat (CC-NB-LRR), (ii) TNL class containing a Toll-interleukin receptor-like domain, a nucleotide binding site and a leucine-rich repeat (TIR-NB-LRR), (iii) RLP class which is a receptor-like protein containing receptor serine-threonine kinase-like domain and an extracellular leucine-rich repeat (ser/thr-LRR), (iv) RLK which are receptor-like kinase containing an extracellular leucine-rich repeat in addition to the kinase domain (Kin-LRR), (v) ‘Others’ class includes any other genes described as using other molecular mechanisms to confer resistance (Sanseverino et al. 2010). Bhatarrai et al. found most of their candidate genes showing high similarity levels to the NB-LRR or LRR gene families and identified two genes as TNL or CNL encoding proteins. Studying potential R genes involved in defense against IDM is paramount in order for breeding IDM-resistant impatiens in the future.

Our research focused on the host side in a compatible reaction between a susceptible cultivar of *I. walleriana* and *P. obducens* during the infection cycle. The main focus was to identify candidate genes that may be involved in impatiens susceptibility to *P. obducens*. These results could provide insights and lay the groundwork for future studies in understanding both host-pathogen interaction in IDM. In this study, we present findings from the first analysis of differential gene expression during the infection of *I. walleriana* by causal agent *P. obducens*.

## Materials and Methods

### *P. obducens* Inoculation and Sample Collection

*Impatiens walleriana* cv. ‘Super Elfin White’ plants were grown in growth chambers maintained at 24°C with a 12 hr light/dark photoperiod. The *P. obducens* isolate was maintained on impatiens plants in the growth chamber. Fully expanded leaves of eight impatiens plants were inoculated by pipetting 50μL sporangial suspension (1×108 sporangia/mL) of *P. obducens* on the abaxial leaf surface. Infection was verified under microscopic observations by staining leaves with tryphan-blue at each time point. Leaves were collected at five different time points (0 hr, 12 hr, 48 hr, 120 hr, and 240 hr) and flash frozen in liquid nitrogen and stored at −80°C until use. Each treatment (time point) consisted of two biological replicates, resulting in a total of 10 leaf tissue RNA samples. Plasmopara obducens sporangia for pathogen baseline samples were carefully scraped off the abaxial surface of impatiens leaves showing sporulation and were also flash frozen and stored before RNA isolation.

### Library Preparation and Sequencing

Total RNA was isolated from *P. obducens* sporangia and inoculated impatiens leaves using the RNeasy Plant Mini Kit (Qiagen, Germantown, MD) according to manufacturer’s protocols. RNA concentration and quality were determined using Bioanalyzer 2100 (Agilent Technologies, Santa Clara, CA). Library sample preparation was done using a stranded mRNA-seq kit (KAPA Biosystems, Inc., Wilmington, MA) and sequencing was performed using an Illumina HiSeq 2500 v4 platform (Illumina, Inc., San Diego, CA) at the Center for Genomic and Computational Biology (GCB) at Duke University. Each library was sequenced using the pair-end (2 × 125bp) protocol. A total of 24 libraries were made (Table 1).

**Table 1:** Statistical summary of RNAseq data.

| Sample ID | Raw Reads (PE) | Clean Reads |
| --- | --- | --- |
| 0H_Control_R1 | 23,520,868 | 18,287,675 |
| 0H_Control_R2 | 27,793,058 | 24,193,950 |
| 12H_Control_R1 | 24,320,452 | 20,299,572 |
| 12H_Control_R2 | 19,672,888 | 15,319,055 |
| 24H_Control_R1 | 24,320,452 | 21,024,513 |
| 24H_Control_R2 | 19,672,888 | 13,922,270 |
| 48H_Control_R1 | 24,162,182 | 17,057,095 |
| 48H_Control_R2 | 17,226,072 | 19,581,661 |
| 120H_Control_R1 | 20,526,634 | 23,639,470 |
| 120H_Control_R2 | 23,321,216 | 18,259,179 |
| 240H_Control_R1 | 27,654,810 | 14,085,034 |
| 240H_Control_R2 | 21,765,004 | 21,712,471 |
| 0H_Infected_R1 | 37,971,712 | 29,005,241 |
| 0H_Infected_R2 | 37,778,302 | 29,248,606 |
| 12H_Infected_R1 | 33,006,920 | 24,968,942 |
| 12H_Infected_R2 | 31,223,426 | 24,440,783 |
| 24H_Infected_R1 | 34,379,442 | 26,500,567 |
| 24H_Infected_R2 | 36,662,544 | 27,860,148 |
| 48H_Infected_R1 | 31,490,402 | 23,140,744 |
| 48H_Infected_R2 | 33,238,084 | 24,795,526 |
| 120H_Infected_R1 | 37,946,602 | 29,078,825 |
| 120H_Infected_R2 | 34,596,692 | 25,346,127 |
| 240H_Infected_R1 | 33,746,888 | 23,867,301 |
| 240H_Infected_R2 | 33,041,348 | 24,552,405 |
| **Total** |  | 528,964,774 |

### RNA-Seq Analysis

Raw reads were imported into CLC Genomics Workbench R9 (Qiagen) and assessed for quality. Sequencing adaptors and low-quality reads were trimmed by 15bp and 5bp from the 5’ and 3’ ends, respectively. A trimming quality score of 0.005 was used. Cleaned high quality reads from samples were assembled de novo using the default parameters in the CLC Genomics Workbench 9.5.3. Because of the complexity of transcriptome data, a word size 25 and bubble size 1000 were used for de novo assembly.

### Assembly Optimization and Functional Annotation

Contigs were subsequently fed to the program cd-hi-est for removal of any redundancy in the sequence, using a sequence identity threshold of 0.95 (Mu□ller et al. 2017). After cd-hit clustering analysis, selected contigs were subjected to comprehensive functional annotation using Blast2GO 5 PRO (Conesa et al. 2005). CloudBLAST tool was used to run BLASTX on transcripts against the non-redundant plant genome NCBI database using an error cutoff value of 1 ×10-5. Gene ontology (GO) mapping of the transcriptome was conducted following annotation of transcripts in order to produce a GO distribution.

### Differential Expression Analysis

Differentially expressed transcripts (DETs) were determined by using the R package DESeq2 using default parameters. Differential expression analysis was done using the transcript counts table generated in CLC genomics. The DETs were filtered at a threshold of log2fold > |2| and padj < 0.001.

A Venn diagram analysis on DETs at all timepoints was done using the online tool provided by VIB and Ghent University (http://bioinformatics.psb.ugent.be/webtools/Venn/). Finally, functions of the DETs, specifically in common plant pathways, were determined using MapMan software.

## Results

### RNA-Seq and de novo Transcriptome Assembly

In total, 689,038,886 reads were generated from all 24 samples (libraries). After quality assessment and trimming, a total of 528,964,774 clean reads were used to construct a de novo assembly in the CLC Genomics workbench which resulted in 91,164 contigs with an N50 value of 1650 bp (50% of transcripts greater than or equal to 1650bp long) (Table 1). These contigs were fed into the cd-hit-est cluster analysis based on similarity, producing 89,987 contigs. Filtering through BLASTx to remove *P. obducens* contigs resulted in 73,022 contigs. This optimized assembly was used as a reference and reads from all the samples were mapped back to this optimized pooled reference assembly and a read count table was generated. Differential expression analysis was done on this table using DEseq2. Only those transcripts were considered as differentially expressed genes that have log2fold > |2| and padj < 0.001. According to this criterion, the number of DETs between infected and control samples ranged from around 3000 to 4500 among infected vs. control samples at all six time points with the highest number of DETs (4549) at 0 hr and the lowest at 120 hr. The number of upregulated genes was more than number of downregulated genes at all time points with the most upregulation happening at 240 hr (2672). Moreover, venn diagram analysis showed that the highest number of DETs (502) unique to timepoint were expressed at 240 hr (Figure 1).

**Figure 1:**
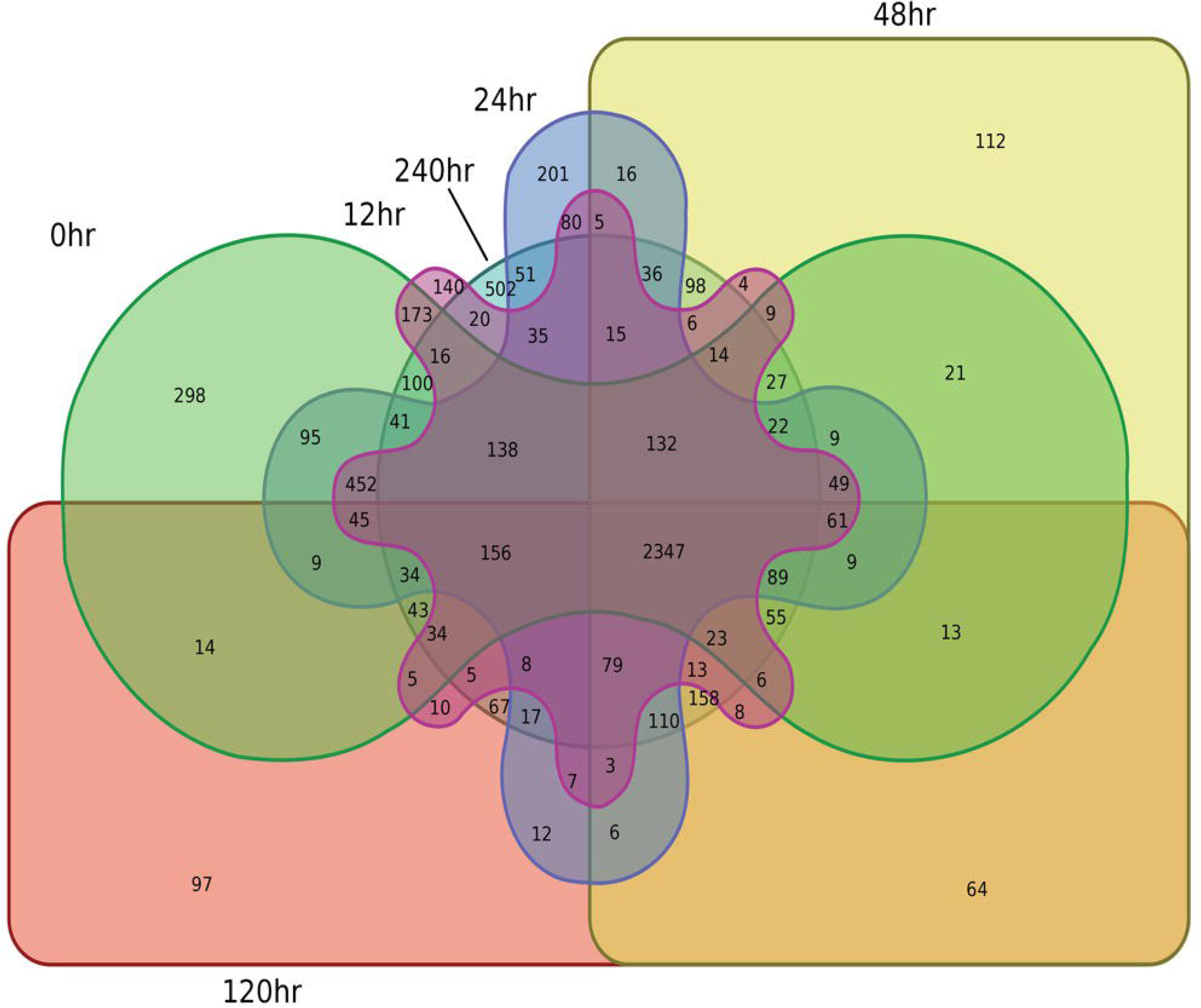
Venn diagram of pair-wise comparisons of DETs unique to timepoints.

### Sequence Annotation

Functional annotation of differentially expressed genes was conducted using Blast2GO software. Species distribution showed little homology with other plant species. There is a lack of impatiens genes available within the NCBI database, therefore the species available with the largest number of BLAST hits was Vitis vinifera (Figure 2). Gene ontology (GO) terms were assigned to DEGS among all three GO categories i.e. biological process (BP), molecular function (MF) and cellular component (CC). Most BP hits were placed in the categories of catalytic and binding and transport activity. The DETs in the MF category were mostly placed within metabolic and cellular processes, response to stimulus and biological regulation. In CC, most DETs fell under the membrane category (Figure 3).

**Figure 2:**
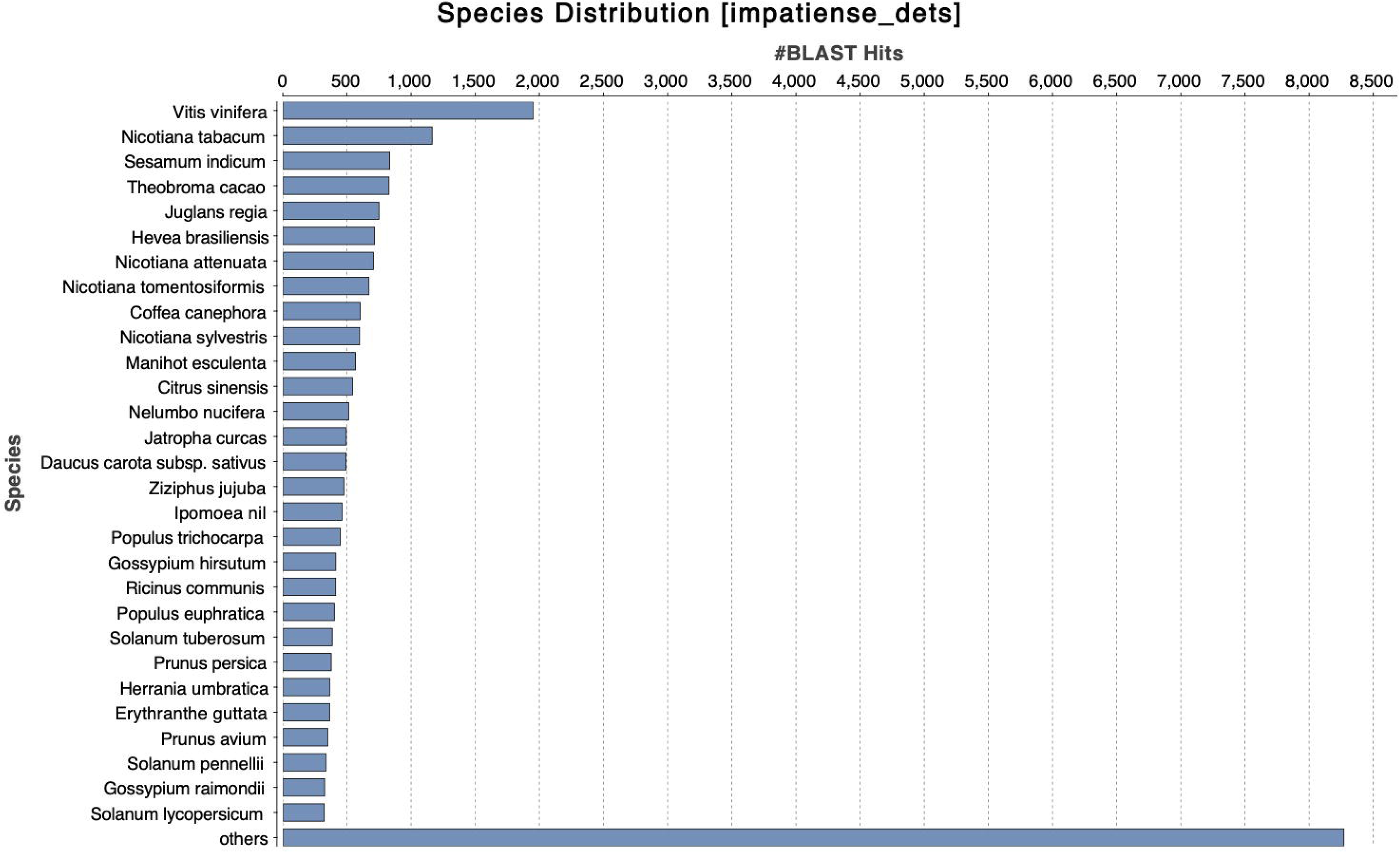
Species distribution showing maximum homology with plant species.

**Figure 3:**
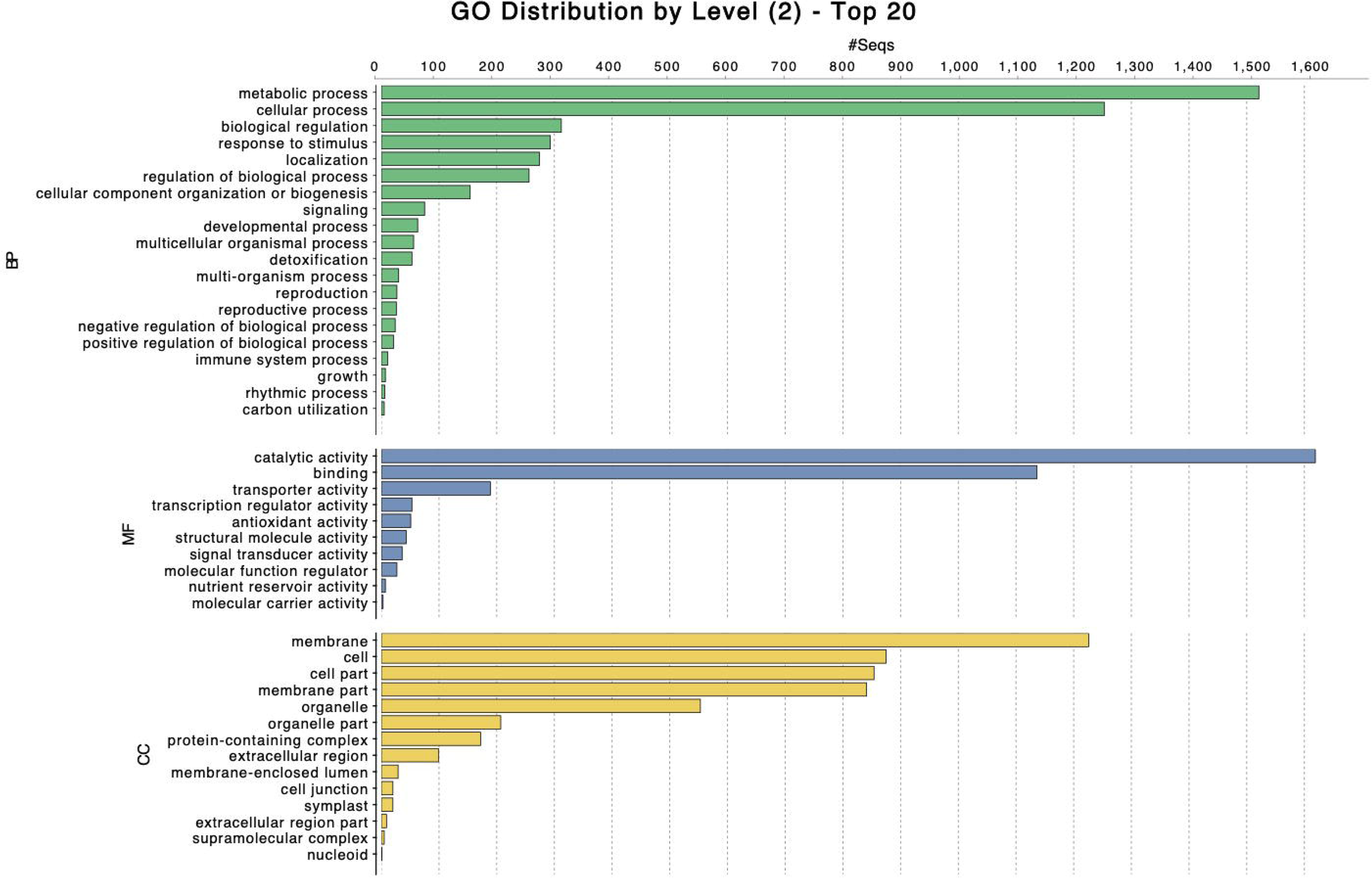
GO sequence distribution chart separated by biological process (BP), molecular function (MF) and cellular component (CC).

We generated MapMan figures at each time point in order to provide a visual representation of genes involved in common plant pathways (Thimm et al. 2004; Usadel et al. 2009). MapMan annotation is only available for few plant species, therefore we used the Arabidopsis homologs for our de novo constructed impatiens transcripts.

DEGs from all six time points were mapped on the biotic stress pathways. Genes involved in proteolysis were observed to be mainly upregulated across all time points (Figures 4). General signaling genes were about equally downregulated as they were upregulated. The WRKY transcription factors (TFs) were mostly upregulated.

**Figure 4:**
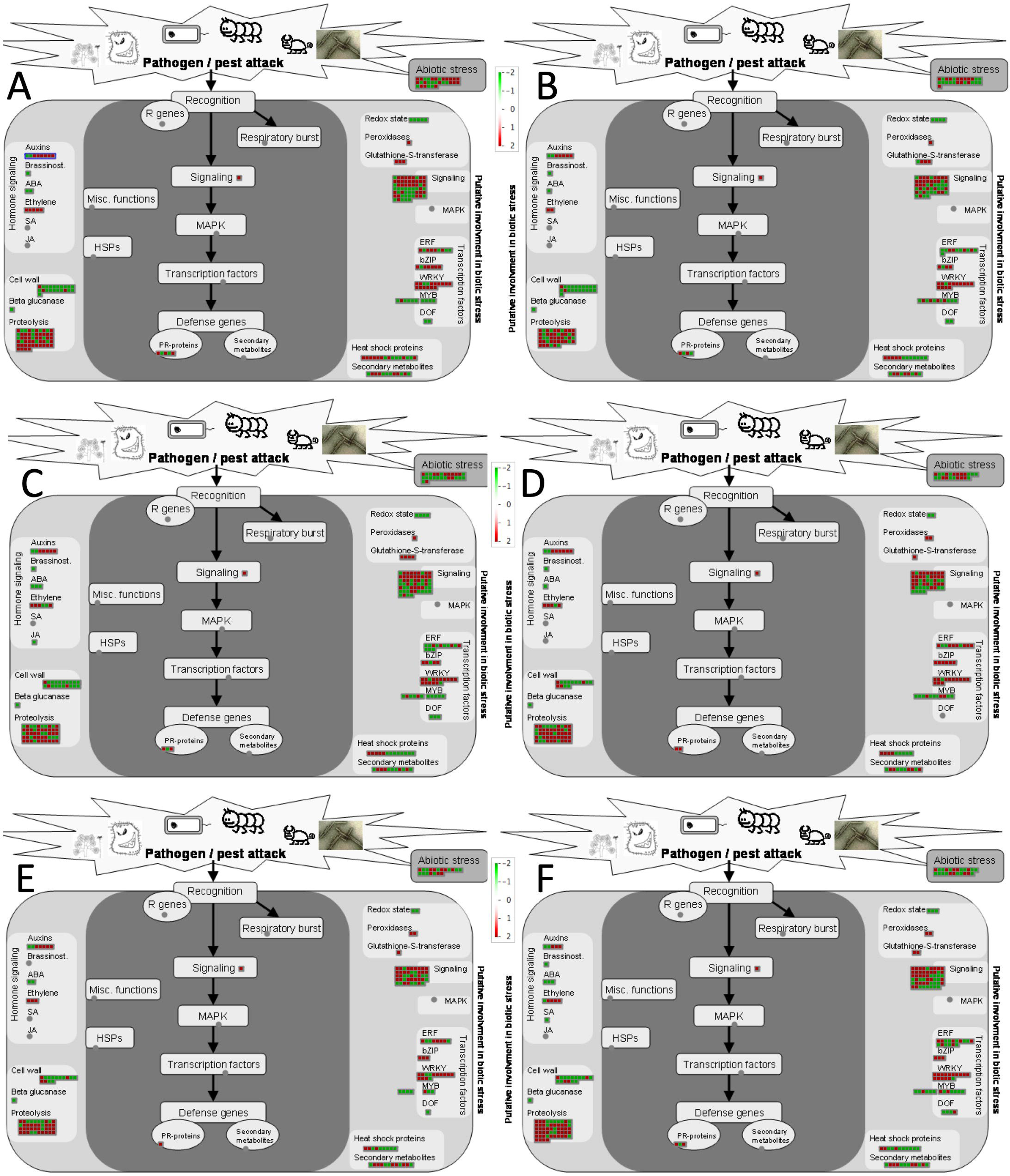
Visualization of differentially expressed genes involved in biotic stress pathways in response to *P. obducens* (A) at 0hpi (hours post-inoculation), (B) at 12hpi, (C) at 24hpi, (D) at 48hpi, (E) at 120hpi and (F) at 240hpi. Color gradient represents log2 fold ratios with red representing upregulation and green representing downregulation in treatments over mock roots. Each box represents one transcript.

### Genes Involved in Plant Defense

Many of already reported plant defense-related genes were found to be differentially expressed at all time points. For example, multiple calcium signaling genes were found differentially expressed at all time points. Most of the calcium binding genes were found to be upregulated, whereas calcium sensing receptors were found to be downregulated across all time points (Table 2).

**Table 2:**
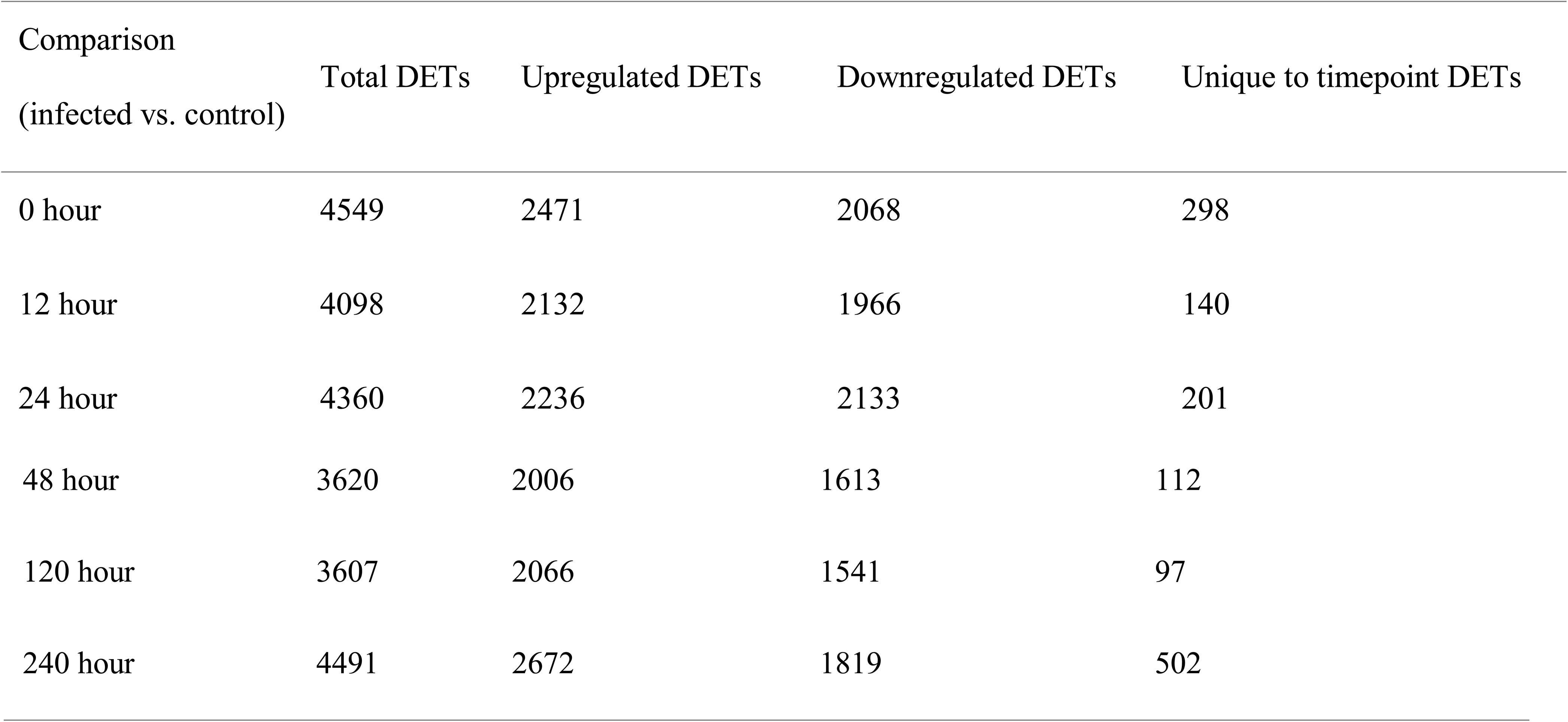
Pairwise comparisons between infected vs. control samples at all six timepoints.

| Comparison<br>(infected vs. control) | Total DETs | Upregulated DETs | Downregulated DETs | Unique to timepoint DETs |
| --- | --- | --- | --- | --- |
| 0 hour | 4549 | 2471 | 2068 | 298 |
| 12 hour | 4098 | 2132 | 1966 | 140 |
| 24 hour | 4360 | 2236 | 2133 | 201 |
| 48 hour | 3620 | 2006 | 1613 | 112 |
| 120 hour | 3607 | 2066 | 1541 | 97 |
| 240 hour | 4491 | 2672 | 1819 | 502 |

Many of the receptor-like kinases (RLKs) were found to be differentially expressed across all time points, including eleven leaf rust 10 disease resistance receptor-like protein kinase transcripts, fifteen G-type lectin S-receptor-like serine/threonine-protein kinase transcripts and five wall-associated receptor kinases. These were found to be mostly upregulated.

We found six different homologs of MLO genes differentially expressed across all time points. Three of these transcripts were upregulated and three were downregulated (Figure 5). Apart from the aforementioned genes, we found many other core plant defense genes differentially expressed in *I. walleriana* in response to infection by *P. obducens* at all stages of the infection cycle.

**Figure 5:**
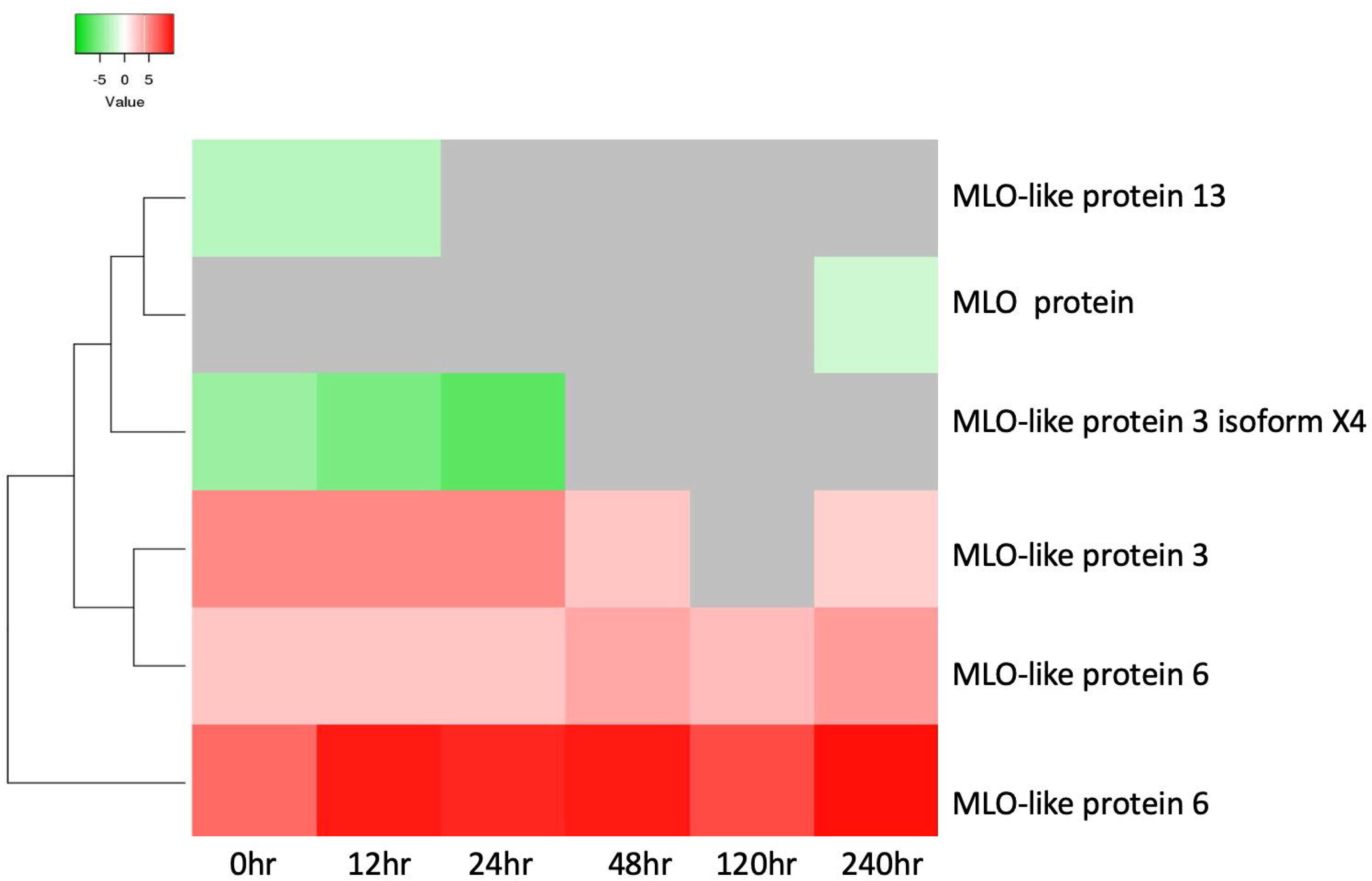
Heat map showing expression profiles of MLO genes.

## Discussion

RNA-Seq is an excellent technique to reveal changes in a plant’s gene expression in response to pathogens, even when whole genome information is unavailable. A recent study was published which is the first comparative transcriptome analysis of both a resistant and susceptible impatiens cultivar in order to identify candidate genes for IDM resistance (Bhattarai et al. 2018). This study was conducted on non-inoculated, uninfected impatiens leaves. To further host-pathogen interaction, we analyzed transcriptional changes in an inoculated, susceptible cultivar of impatiens at 6 timepoints. At 10 days (240 hr) post-inoculation, sporulation of *P. obducens* can be observed on the abaxial leaf surface of susceptible, infected impatiens plants. *Plasmopara obducens* is an obligate biotroph, therefore we wanted to track transcriptional changes throughout the entirety of its infection cycle (0 to 240 hr). To our knowledge, this is the first study of the host-pathogen interaction between *I. walleriana* and *P. obducens* available.

In 2018, Ball Horticultural Company and KeyGene announced the completion of the entire genome sequencing and assembly of *I. walleriana*. Because it is not currently available to the public, our study relied on the use of de novo transcriptome assembly of *I. walleriana*. After assembly, optimization was done in order to preserve biologically meaningful transcripts for further differential expression analysis. Differential gene expression among infected and control samples were observed immediately after inoculation. In order to track transcriptional changes during the first contact between plant and pathogen, we collected samples for the first time point immediately following inoculation, designated as 0hr. Surprisingly, we observed the highest number of differentially expressed genes at 0hr than at any other timepoint, except 240 hr.

Plant pathogens contain specific structures or molecules known as pathogen associated molecular patterns (PAMP), which are recognized by PRRs to active plant defense. This is known as PAMP-Triggered Immunity (PTI). Once activated, PTI results in basal defense responses to pathogen invasion (Dodds and Rathjen 2010). The present study showed the differential expression of all kinds of PRRs in susceptible impatiens infected with *P. obducens* at 0hr. Plants carry two types of PRRs: receptor-like kinases (RLKs) and receptor-like proteins (RLPs) (Tang et al. 2017). Wall-associated RLKs have been reported as the positive regulators of plant defense against fungi (Delteil et al. 2016). In our study, we found five wall-associated receptor-like kinases to be upregulated.

Another type of RLK, G-type lectin S-receptor-like serine/threonine-protein kinase, was also found to be upregulated across all timepoints, including 0 hr. These lectin receptor kinases are a class of RLKs divided into three subclasses (C-type, G-type and L-type). The G-type kinase that we have found upregulated in this study has yet to be reported in oomycete-plant interactions. Further studies may be able to confirm if G-type receptor kinases play a role in plant defense against oomycetes. We also found eleven leaf rust 10 disease resistance receptor-like protein kinase isoforms to be upregulated as well. This is an indication that the impatiens plant recognizes infection by *P. obducens* immediately upon inoculation, prompting plant defenses. Membrane-associated PRRs have previously been reported to be involved in initial plant-pathogen interactions, but we observed the continuation of upregulation of these RLKs throughout the entire infection cycle of *P. obducens* in *I. walleriana*.

After recognition by PRRs, calcium ion (Ca2+) concentrations are considered to be the next event in plant responses to any environmental cues (Ranty et al. 2016). Many calcium signaling genes in this study were found to be differentially expressed starting at initial contact between *P. obducens* and *I. walleriana*. Our study focused on a compatible interaction between a susceptible cultivar of *I. walleriana* and *P. obducens*. Many transcriptome studies have revealed there is involvement of Ca2+ signaling genes in compatible host-pathogen interactions (Aldon et al. 2018; Naveed and Ali 2018).

MLO proteins were initially identified as common powdery mildew susceptibility genes, but recent studies have found them to be involved in plant-oomycete interactions as well (Zhu et al. 2017; Naveed and Ali 2018). We found six homologs of MLO genes to be differentially expressed in *I. walleriana*, three of which were highly upregulated. Induction of MLO genes by *P. obducens* in *I. walleriana* could be speculated as another example of MLO genes being involved in host susceptibility to pathogens other than powdery mildew, in this case, an oomycete.

In addition to the aforementioned genes, we found many other core plant defense-related genes such as pathogenicity-related genes, resistance genes, and biotic stress-related transcription factors to be differentially expressed throughout the infection cycle of IDM. This study revealed the presence of transcriptional changes in common physiological and core plant defense-related pathways in *I. walleriana* during infection by *P. obducens*. Many candidate genes have been identified in this study,

## References

Aldon, D., Mbengue, M., Mazars, C., and Galaud, J.-P. 2018. Calcium signaling in plant biotic interactions. Int. J. Mol. Sci. 19: 665.

Bhattarai, K., Wang, W., Cao, Z. and Deng, Z. 2018. Comparative analysis of impatiens leaf transcriptomes reveal candidate genes for resistance to downy mildew caused by Plasmopara obducens. Int. J. Mol. Sci. 19: 2057.

Delteil, A., Gobbato, E., Cayrol, B., Estevan, J., Michel-Romiti, C., Dievart, A., Kroj, T., and Morel, J. B. 2016. Several wall-associated kinases participate positively and negatively in basal defense against rice blast fungus. BMC Plant Biology. 16:17.

Dodds, P. N., and Rathjen, J. P. 2010. Plant immunity: towards an integrated view of plant-pathogen interactions. Nature Reviews Genetics. 11: 539–548.

Farr, D. F., and Rossman, A. Y. 2016. Fungal Databases, Systematic Mycology and Microbiology Laboratory, ARS, USDA.

Lane, C. R., Beales, P. A., O’Neill, T. M., McPherson, G. M., Finlay, A. R., David, J., Constantinescu, O., and Henricot, B. 2005. First report of Impatiens downy mildew (Plasmopara obducens) in the UK. Plant Pathol. 54:243.

Naveed, Z. A., and Ali, G. S. 2018. Comparative transcriptome analysis between a resistant and a susceptible wild tomato accession in response to Phytophthora parasitica. Int. J. Mol. Sci. 19: 3735.

Salgado-Salazar, C., Rivera, Y., Veltri, D., and Crouch, J. A. 2015. Polymorphic SSR markers for Plamopara obducens (Peronosporaceae), the newly emergent downy mildew pathogen of Impatiens (Balsaminaceae). Appl. Plant. Sci. 3: 1500073.

Sanseverino, W., Roma, G., De Simone, M., Faino, L., Melito, S., Stupka, E., Frusciante, L., and Ercolano, M.R. 2010. Nucleic Acids Res. 38: D814–D821.

Tang, D., Wang, G., and Zhou, J. 2017. Receptor kinases in plant-pathogen interactions: more than pattern recognition. Plant Cell. 29: 618–637.

Thimm, O., Blasing, O., Gibon, Y., Nagel, A., Meyer, S., Kruer, P., Selbig, J., Muller, L. A., Rhee, S. Y., and Stitt, M. 2004. Mapman: a user driven tool to display genomics data sets onto diagrams of metabolic pathways and other biological processes. The Plant Journal. 37: 914–939.

Zhu, Y., Shao, J., Zhou, Z., and Davis, R.E. 2017. Comparative transcriptome analysis reveals a preformed defense system in apple root of a resistant genotype of G.935 in the absence of pathogen. Int. J. Plant Genomics. 2017: 895076.

